# A cell-autonomous PD-1/PD-L1 circuit promotes tumorigenicity of thyroid cancer cells by activating a SHP2/Ras/MAPK signalling cascade

**DOI:** 10.1101/2020.09.22.307942

**Authors:** Federica Liotti, Narender Kumar, Nella Prevete, Maria Marotta, Daniela Sorriento, Caterina Ieranò, Andrea Ronchi, Federica Zito Marino, Sonia Moretti, Renato Colella, Efiso Puxeddu, Simona Paladino, Yoshihito Kano, Michael Ohh, Stefania Scala, Rosa Marina Melillo

## Abstract

The programmed cell death-1 (PD-1) and its ligands PD-L1 and PD-L2 are immune checkpoints. Typically, cancer cells express the PD-Ls that bind PD-1 on immune cells, inhibiting their anti-cancer activity. Recently, PD-1 expression has been found in cancer cells. We analysed expression and functions of PD-1 in thyroid cancer (TC). Human TC specimens (47%), but not normal thyroids, displayed PD-1 expression in epithelial cells, which significantly correlated with tumour stage and lymph-node metastasis. PD-1 overexpression/stimulation promoted TC cell proliferation and migration in culture. PD-1 recruited the SHP2 phosphatase, potentiated its phosphatase activity thus enhancing Ras activation by dephosphorylation of inhibitory tyrosine 32 and triggering the MAPK cascade. PD-1 inhibition decreased, while PD-1 overexpression facilitated, TC cell xenograft growth by affecting cell proliferation. PD-1 circuit blockade in TC, besides restoring anti-cancer immunity, could also directly impair TC cell growth by inhibiting the Ras/MAPK pathway.

## Introduction

Immunotherapy represents the major breakthrough of the last years in the therapy of several cancer types (*Yang, 2015*). The programmed cell death-ligand 1 and 2 (PD-L1, PD-L2) are immune checkpoints (IC) important for delivering inhibitory signals to immune cells expressing their receptor programmed cell death-1 (PD-1) (*Yang, 2015*). This circuit is critical in regulating immune tolerance in various physiologic and pathologic contexts (*Yang, 2015*). Cancer cells suppress anti-cancer immune response exploiting the PD-1 circuit (*Rabinovich et al., 2007*). Typically, PD-Ls are expressed by cancer cells, while PD-1 is expressed by immune cells with anti-cancer potential (i.e., T cells, macrophages or natural killer cells) (*Rabinovich et al., 2007*). The inhibition of this circuit through immune checkpoint inhibitors (ICI) - neutralizing antibodies against PD-1, PD-L1 or PD- L2 - restores the anti-cancer immune response and displays therapeutic activity in various cancer types (*McNutt, 2013*).

Recently, various tumour types have been found to express also intrinsic PD-1 (i.e., melanoma, hepatocarcinoma, lung carcinoma and T-cell lymphomas) (*Kleffel et al., 2015, Li et al., 2017, Du et al., 2018, Zhao et al., 2018*). PD-1 intrinsic signalling promoted tumour growth in melanoma and hepatocarcinoma through a mammalian target of rapamycin (mTOR)/ribosomal protein S6 Kinase (S6K1) pathway (*Kleffel et al., 2015, Li et al., 2017*). By contrast, in non-small cell lung cancer (NSCLC) and in T-cell lymphomas, PD-1 behaved as a tumour suppressor (*Du et al., 2018, Zhao et al., 2018*). These data indicate that PD-1 could exert context-related tumour-intrinsic functions other than the suppression of immune response, and suggest the need of wider studies on ICI effects on the entire tumour context.

Thyroid carcinoma (TC) is the most frequent endocrine malignancy. Follicular cell-derived TC includes different histotypes ranging from well differentiated (WDTC) to poorly differentiated (PDTC) and undifferentiated/anaplastic (ATC) carcinomas. WDTCs include papillary histotype (PTC), representing the majority of these tumours, and follicular histotype (FTC). WDTCs show an indolent behaviour and are mainly cured by surgery and ^131^I radioiodine (RAI) therapy; only a small percentage of them exhibits recurrence, metastasis and resistance to RAI over time. By contrast, aggressive forms of TC (PDTC and ATC) represent a clinic challenge displaying a remarkable chemo- and radio-resistant phenotype from the beginning (*Naoum et al., 2018, Liotti et al., 2019*). Interestingly, aggressive forms of TC exhibit increased immune checkpoint expression and inefficient immune infiltrate (*French et al., 2012, Bastman et al., 2016, Giannini et al., 2019, Liotti et al., 2019, Malfitano et al., 2019*), features that are being evaluated for the treatment of the disease (*Saini et al., 2018, Liotti et al., 2019, Malfitano et al., 2019*).

Here, we analysed the PD-1/PD-Ls circuit in TC showing that: i) TC cell lines and TC human samples express, besides PD-Ls, as already demonstrated (*Cunha et al., 2012, Cunha et al., 2013, Ulisse et al., 2019*), also PD-1 at epithelial level, whose levels correlated with tumour aggressiveness; ii) intrinsic PD-1 sustains proliferation and migration of TC cells through a SHP2/Ras/MAPK signalling cascade; iii) PD-1 overexpression promotes, while PD-1 blockade inhibits, ATC xenograft growth by affecting cancer cell proliferation.

Thus, TCs express an intrinsic pro-tumorigenic PD-1 circuit. In TC context, the oncogenic role of PD-1 is dependent on the activation of the Ras/MAPK cascade. PD-1 blockade may represent a rational therapeutic choice in aggressive forms of TC for both immune response reconstitution and direct anti-tumour effects.

## Results

### PD-1 receptor and its ligands are expressed in thyroid carcinoma cells

We evaluated the expression levels of PD-1, PD-L1 and PD-L2 in a panel of human TC cell lines derived from PTC (BcPAP, TPC-1) or ATC (8505c, CAL62, SW1736, FRO, BHT101, HTH7, OCUT1) compared to a primary human thyroid cell culture (H-6040). Cytofluorimetric analysis demonstrated that all the cell lines expressed PD-1 on the plasma membrane, though to a lesser extent than PD-Ls, and that PD-1 protein levels were higher in cancer compared to normal thyroid cells (**Figure 1A**). PD-1, PD-L1 and PD-L2 mRNA levels were comparable between normal and cancerous thyroid cells, suggesting that post-translational mechanisms could be responsible for the protein increase observed in cancer cells (**Figure Supplement 1**).

**Figure 1.**
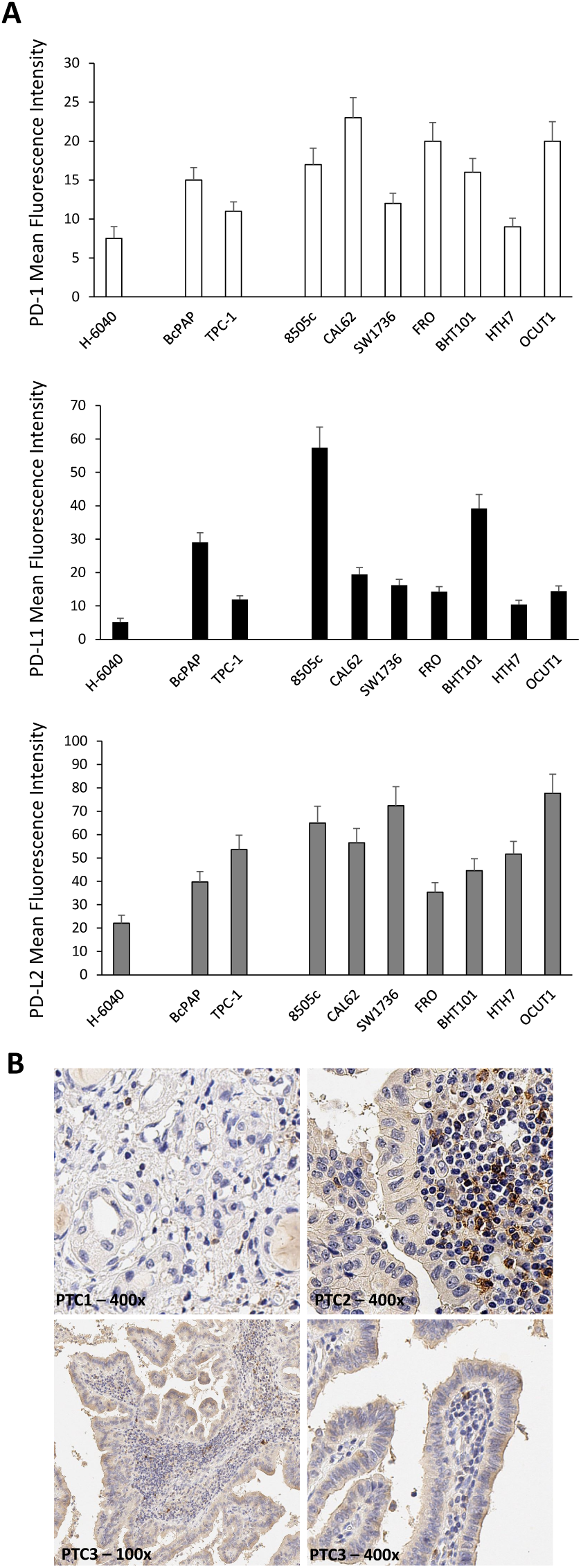
Immune checkpoint expression in thyroid cancer (TC) cells and human TC tissue samples. **A**. Mean Fluorescence Intensity for PD-1, PD-L1 and PD-L2 measured by flow cytometric analysis on H-6040 normal thyroid epithelial cells, PTC-derived cell lines (BcPAP and TPC-1), and ATC-derived cell lines (8505c, CAL62, SW1736, FRO, BHT101, HTH7, OCUT1). Data are presented as mean ± SD. **B**. Immunohistochemical staining of representative PTC samples with anti-PD-1 antibody. Examples of negative PD-1 staining (**PTC1**), intense PD-1 staining in the tumour immune infiltrate (**PTC2**), and PD-1 immunoreactivity in thyroid epithelial cells (**PTC3**) are shown. In positive samples, immunoreactivity was detected mainly in the cytosolic portion and/or at plasma membrane of epithelial TC cells.

Immunohistochemical (IHC) staining of whole sections from 34 PTC surgical samples with anti-PD-1 antibodies showed that PD-1 is expressed in TC cells (**Figure 1B**), but not in normal thyroid epithelial cells (**not shown**). **Figure 1B** shows a representative PTC case with negative PD-1 staining (**PTC1**), intense PD-1 staining in the tumour immune infiltrate (**PTC2**), and PD-1 immunoreactivity, cytosolic and/or localized at the plasma membrane, in thyroid cancer epithelial cells (**PTC3**).

PD-1 expression was detectable in epithelial cancerous cells of 47% of tumour samples (**Table 1**). By analysing clinic-pathologic features of the PTC samples, we found that tumour stage and lymph-nodal metastasis significantly correlated with PD-1 staining (**Table 1**) in our casistic.

**Table 1.** Relation between PD-1 epithelial TC cell expression and clinic-pathological features. * P < 0.05 among groups

|  | Epithelial cell PD-1 staining |  |  |
| --- | --- | --- | --- |
|  | Negative/low | Positive | P |
| **BRAF (n=34)** |  |  |  |
| not mutated | 2 | 2 | 0.9 |
| mutated | 16 | 14 |  |
| **Thyroiditis (n=33)** |  |  |  |
| No | 12 | 11 | 0.9 |
| Yes | 5 | 5 |  |
| **TNM (n=34)** |  |  |  |
| T1 | 11 | 9 | 0.77 |
| T2 | 4 | 3 |  |
| T3 | 3 | 4 |  |
| N0 | 17 | 6 | **0.04\*** |
| N1 | 1 | 10 |  |
| M0 | 11 | 15 | 0.9 |
| M1 | 7 | 1 |  |
| **Progression free survival (n=28)** |  |  |  |
| No | 3 | 4 | 0.51 |
| Yes | 12 | 9 |  |
| **Stage (n=29)** |  |  |  |
| 1 | 12 | 6 | **0.04\*** |
| 2 | 0 | 0 |  |
| 3 | 1 | 5 |  |
| 4 | 2 | 3 |  |
| **Infiltrative phenotype (n=34)** |  |  |  |
| No | 15 | 13 | 0.87 |
| Yes | 3 | 3 |  |

These data indicate that TC cells can express PD-1 together with its ligands (*Cunha et al., 2012, Cunha et al., 2013, Ulisse et al., 2019*), and that PD-1 expression correlates with tumour malignancy.

### PD-1 promotes thyroid carcinoma cell proliferation and motility

We selected 8505c and TPC-1 cells - derived from a human ATC and PTC, respectively - to analyse the biologic effects of PD-1 enforced expression or of PD-1 stimulation by soluble PD-L1 (sPD-L1 - 1 μg/ml). Endogenous PD-1 protein expression levels in these cell lines, together with levels of PD-1 expression upon transient transfection, is shown in **Figure Supplement 2A**. We demonstrated that transient PD-1 overexpression (pFLAG PD-1 compared to pFLAG) or PD-1 activation (sPD-L1 *vs* untreated - NT) significantly increased DNA synthesis, as assessed by BrdU incorporation (**Figure Supplement 2A**) in both TC cell lines. Accordingly, cell cycle analysis showed an increased percentage of cells in S and G2/M phases in PD-1-transfected compared to empty vector-transfected TC cells (**Figure Supplement 2B**). No effects of PD-1 overexpression/activation were observed on cell viability (**Figure Supplement 2C**). In order to confirm these observations, we evaluated the effects of PD-1 inhibition on the same cellular functions. To this aim, PD-1 expression was inhibited by siRNA or Nivolumab (anti-PD-1 moAb) in TPC-1 and 8505c cells.

Both siRNAs targeting PD-1 (siPD-1 *vs* siCTR - 100 nM; **Figure Supplement 2D**) and Nivolumab (10 μg/ml) (Nivo *vs* IgG_4_) were able to significantly inhibit BrdU incorporation (**Figure 2B**) and cell cycle progression (**Figure Supplement 2E**) of TC cells in comparison to the relative controls, without affecting cell viability (**Figure Supplement 2F**).

**Figure 2.**
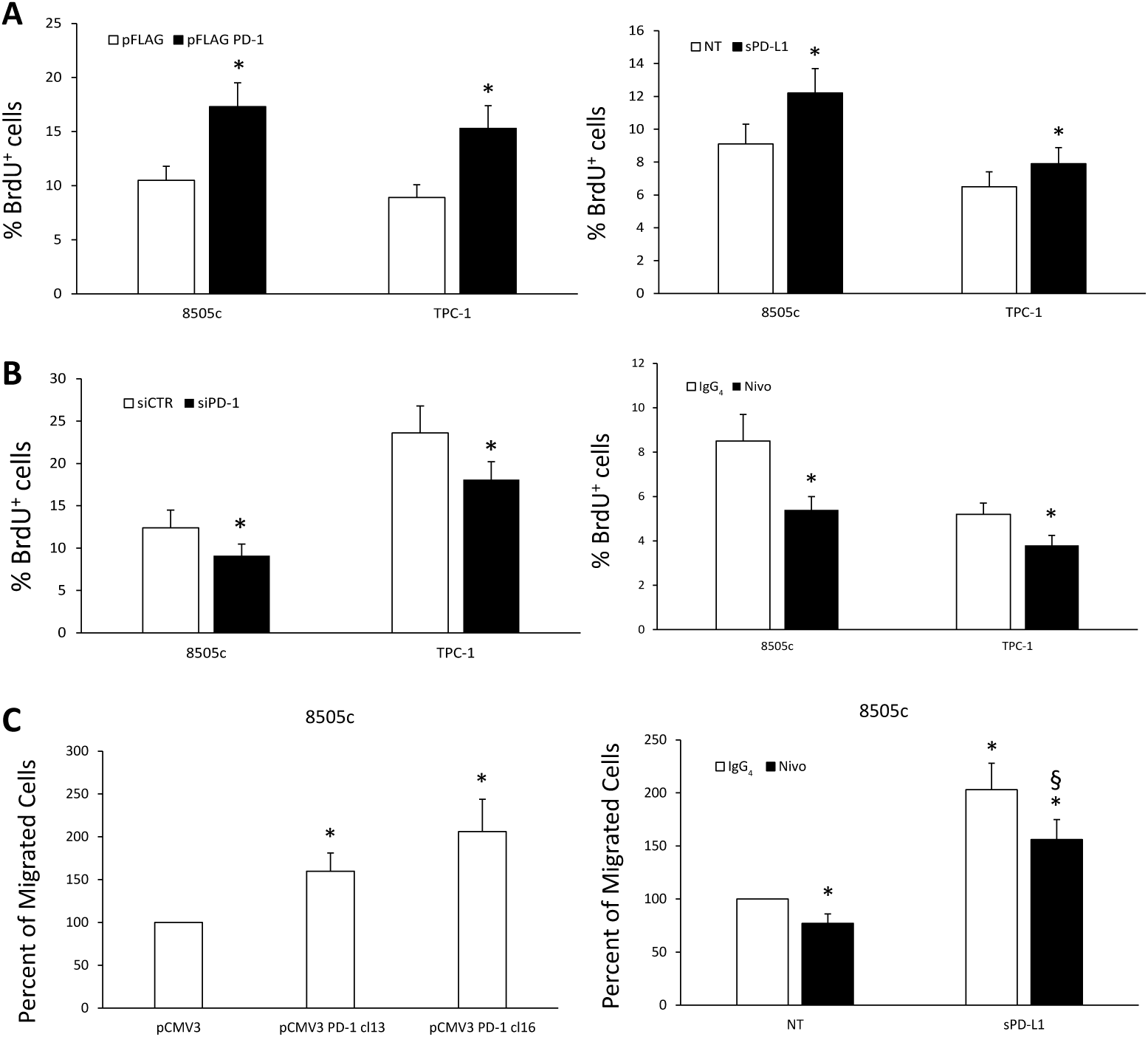
Functional activity of intrinsic PD-1 in TC cells. **A**. DNA synthesis of 8505c and TPC-1 cells transiently transfected with pFLAG PD-1 or the relative empty vector (pFLAG), or treated or not with soluble PD-L1 (sPD-L1 – 1 μg/ml) assessed by BrdU incorporation. Data are presented as mean ± SD of 5 independent experiments. **B**. DNA synthesis of 8505c and TPC-1 cells treated with siRNA targeting PD-1 (siPD-1 – 100 nM) or the relative control (siCTR – 100 nM) for 48 h or with Nivolumab (10 μg/ml) of control IgG_4_ (10 μg/ml) for 24 h assessed by BrdU incorporation. Data are presented as mean ± SD of 5 independent experiments. **C**. Percent of migrated cells over control of stably transfected 8505c PD-1 cells versus 10% FBS, or 8505c cells treated with Nivolumab (10 μg/ml) or control IgG_4_ (10 μg/ml) toward sPD-L1 (1 μg/ml) or medium alone. Data are presented as mean ± SD of 5 independent experiments. \**P*<0.05 compared to the relative untreated cells. § *P*<0.05 compared to sPD-L1 alone.

To assess the role of endogenous PD-1 ligands in TC cell proliferation, we treated 8505c cells with blocking anti-PD-L1 or anti-PD-L2 moAb (10 μg/ml) or transiently transfected them with PD-L1 or PD-L2 expressing vectors. PD-L1 or PD-L2 overexpression increased, while anti-PD-L1 or anti-PD-L2 antibodies inhibited, BrdU incorporation in 8505c cells (**Figure Supplement 2G**). No effects of PD-L1 or PD-L2 were observed on TC cell viability (**not shown**).

Since PD-1 expression levels in human TC samples correlated with lymph-nodal metastasis, we asked whether PD-1 could also stimulate the motility of TC cells. To this aim, we performed migration assays on 8505c cells stably overexpressing or not PD-1 [pCMV3 PD-1 cl13 and cl16 compared to pCMV3 empty vector-transfected cells (**Figure Supplement 3A**)] or on parental 8505c cells treated or not with sPD-L1 (1 μg/ml) in the presence or absence of Nivolumab (10 μg/ml) (**Figure 2C**). PD-1 overexpressing TC cells showed increased migratory potential compared to control cells. Consistently, sPD-L1 induced, and Nivolumab inhibited, both basal and sPD-L1-induced migration (**Figure 2C**).

These data indicate that PD-1 intrinsic circuit sustains TC cell proliferation and migration.

### PD-1 activates the Ras/MAPK signalling cascade in thyroid carcinoma cells

We then asked which signalling pathway was stimulated upon PD-1 overexpression/activation. To this aim, we used specific phospho-antibodies against various signalling proteins. We found that BRAF, MEK and MAPK (p44/p42) are activated, as demonstrated by increased levels of their phosphorylated forms, upon PD-1 transient transfection (**Figure 3A**), PD-1 stable transfection (**Figure Supplement 3B**), and sPD-L1 treatment (**Figure 3B**) in both 8505c and TPC-1 cells. No significant activation of other signalling proteins was detected (**Figure Supplement 3C**). To confirm these observations, BRAF, MEK1/2 and MAPK activation levels were evaluated upon PD-1 blockade by siPD-1 or Nivolumab treatment. Consistently, both siPD-1 (100 nM) and Nivolumab (10 μg/ml – 15 and 30 min) reduced the levels of phosphorylated BRAF, MEK1/2 and MAPK compared to the relative controls (**Figure 3C**) in TC cells.

**Figure 3.**
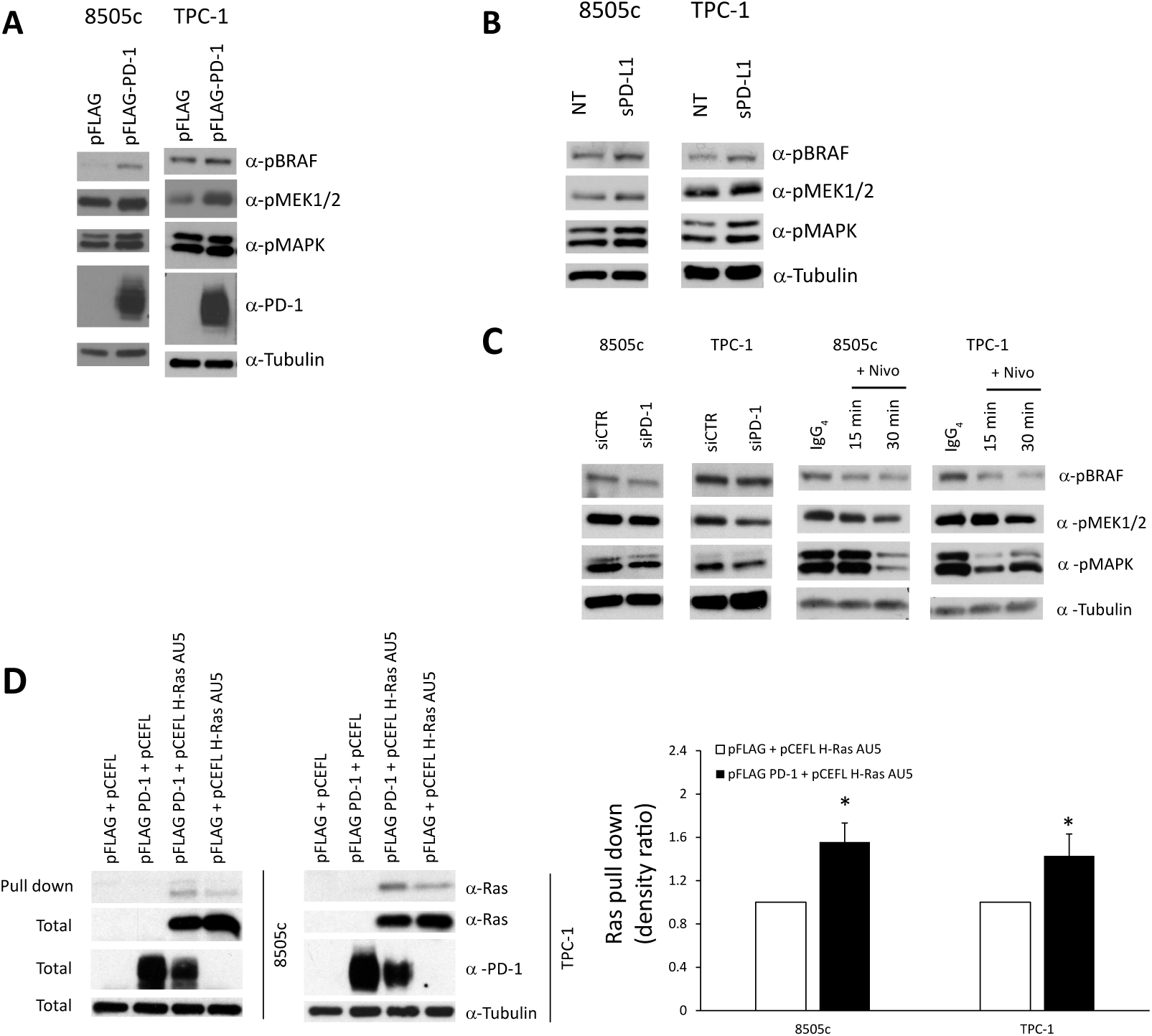
Signalling pathways downstream PD-1 activation/overexpression. **A**. Activation of BRAF, MEK1/2 and MAPK (p44/p42) in 8505c and TPC-1 cells, transfected with pFLAG PD-1 or the relative empty vector (pFLAG), assessed by western blot for their phosphorylated forms. **B**. Activation of BRAF, MEK1/2 and MAPK in 8505c and TPC-1 cells, treated or not with sPD-L1 (1 μg/ml - 30 min), assessed by western blot for their phosphorylated forms. **C**. Activation of BRAF, MEK1/2 and MAPK in 8505c and TPC-1 cells, treated with siPD1 or siCTR (100 nM - 48 h) or with Nivolumab or IgG_4_ (10 μg/ml – 15 and 30 min), assessed by western blot for their phosphorylated forms. **D**. Pull-down assay with the GST-RAF1-Ras Binding Domain (RBD) of 8505c and TPC-1 cells transiently transfected with pFLAG + pCEFL, pFLAG PD-1 + pCEFL, pFLAG + pCEFL H-Ras AU5, or pFLAG PD-1 + pCEFL H-Ras AU5. A representative pull-down is shown, together with the mean densitometric analysis ± SD of 5 independent assays. * *P*<0.05 compared to the relative control.

Since the BRAF/MEK/MAPK signalling is potentiated by PD-1 in TC cells, and Ras GTPase is the main upstream activator of this cascade (*Knauf and Fagin, 2009*), we asked whether PD-1 could activate Ras. To this end, we used a pull-down assay with the GST-RAF1-Ras binding domain (RBD), which specifically binds the GTP-loaded active form of Ras. 8505c and TPC-1 cells were transiently transfected with empty vector (pFLAG) or PD-1 (pFLAG PD-1) in combination with pCEFL H-Ras AU5 or the relative empty vector (pCEFL). PD-1 enforced expression increased Ras activation, as assessed by Ras pull-down, in comparison to control (**Figure 3D)**, suggesting that PD-1 potentiates Ras activation in TC cells.

### PD-1 recruits and activates the SHP2 phosphatase in thyroid carcinoma cells

In immune cells, PD-1 signalling requires the tyrosine phosphatase SHP2 (PTPN11) (*Bunda et al., 2015*). Upon phosphorylation of tyrosine residues in its cytosolic domain, PD-1 binds to the SH2 domains of SHP2 that, in turn, dephosphorylates signalling components of the immune receptors, thus down-regulating the immune responses (*Rota et al., 2018*). In cancer cells, SHP2 acts as a signalling molecule downstream receptor tyrosine kinases (RTKs), displaying oncogenic activity (*Zhang et al., 2015*). In particular, SHP2 can contribute to Ras activation either by recruiting the GRB2/SOS complex to the plasma membrane (*Ran et al., 2016*) or through its phosphatase activity on Ras inhibitory tyrosine residues (*Matozaki et al., 2009, Ran et al., 2016*).

We first asked whether PD-1 could physically interact with SHP2 in TC cells. Reciprocal co-immunoprecipitation experiments showed that endogenous and exogenously expressed PD-1 bind SHP2 in 8505c and TPC-1 cells (**Figure 4A**). Moreover, pull-down assays with N- or C-terminal SH2 domain of SHP2 demonstrated that SHP2 can bind PD-1 mainly through SHP2 C-terminal SH2 domain (**Figure 4B**). In support of these observations, we found that both endogenous and exogenous PD-1 are tyrosine phosphorylated in TC cells (**Figure Supplement 4A**), condition necessary to allow the SH2 domains of SHP2 to bind PD-1 (*Ran et al., 2016*).

**Figure 4.**
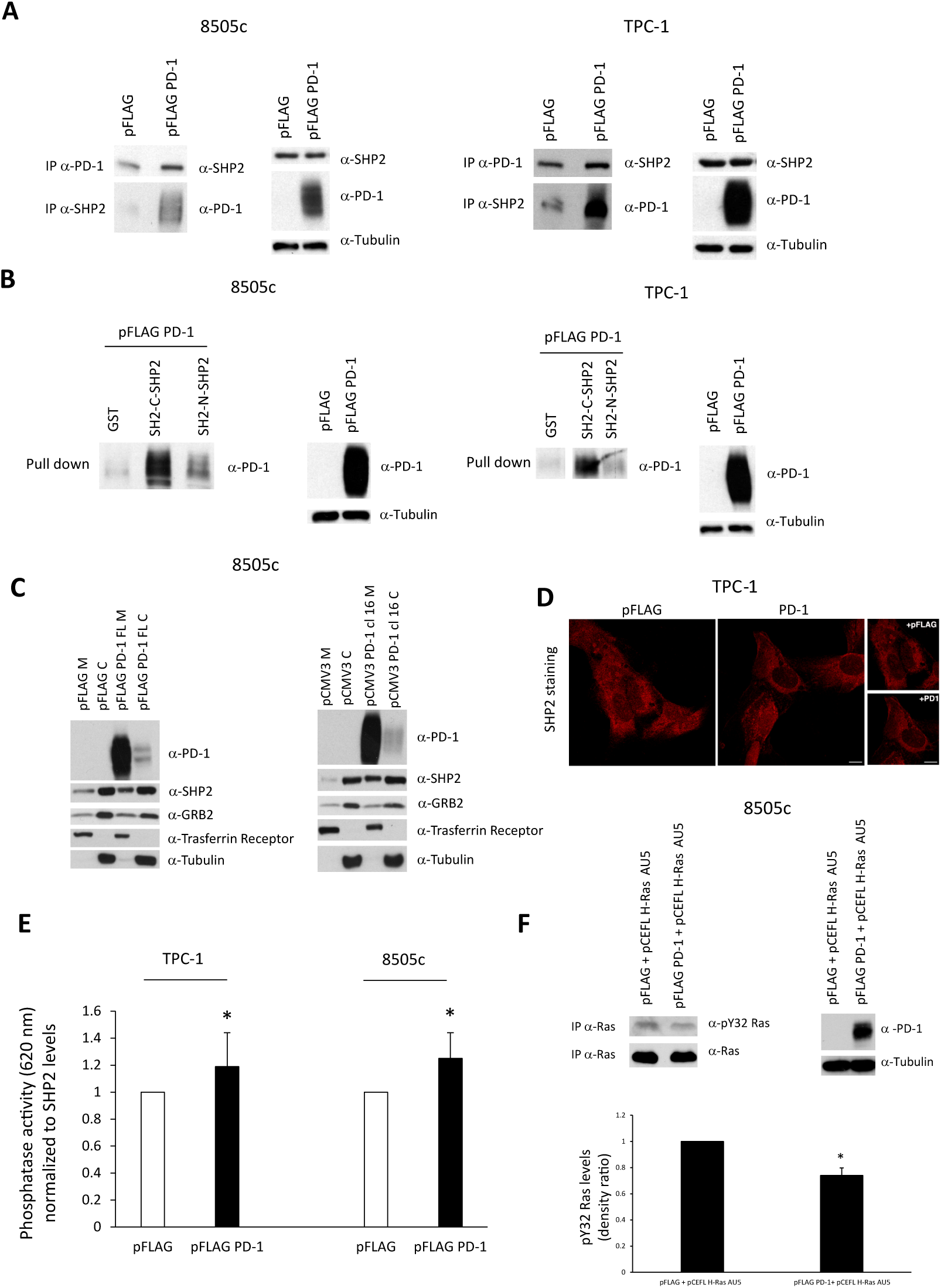
Effects of intrinsic PD-1 on SHP2 localization and functions. **A**. Total cell protein extracts from 8505c and TPC-1 cells transiently transfected with pFLAG PD-1 or the empty vector (pFLAG) were subjected to immunoprecipitation followed by western blotting with the indicated antibodies. **B**. Total protein extracts from 8505c and TPC-1 cells transiently transfected with pFLAG-PD-1 were subjected to an *in vitro* pull-down assay using the indicated recombinant proteins. Bound proteins were immunoblotted with antibody against PD-1. **C**. 8505c cells transiently transfected with PD-1 (pFLAG PD-1) or stably overexpressing PD-1 (pCMV3 PD-1 cl16) and the relative control cells were harvested and subjected to cell protein fractionation. Membrane (M) and cytoplasmic (C) protein fractions were immunoblotted with the indicated antibodies. Transferrin receptor or tubulin levels were used as normalization of membrane and cytosolic fractions, respectively. **D**. Immunofluorescence microscopy of TPC-1 cells, transiently transfected with pFLAG PD-1 or the empty vector, stained with an antibody specific for SHP2. Bars, 5 μm. **E**. SHP2 phosphatase activity assay on TPC-1 and 8505c cells transiently transfected with PD-1 (pFLAG PD-1) and the relative control (pFLAG), assessed by using a specific SHP2 phosphorylated substrate in the presence of the Malachite Green tracer, a colorimetric method (absorbance at 620 nm) for the detection of free inorganic phosphate. The SHP2 phosphatase activity was normalized for SHP2 content as assessed by western blot. Data are presented as mean ± SD of 3 independent experiments. **F**. Total cell protein extracts from 8505c cells transiently transfected with pCEFL H-Ras AU5 + pFLAG PD-1 or empty vector (pFLAG) were subjected to immunoprecipitation followed by western blotting with the indicated antibodies. A representative experiment is shown, together with the mean densitometric analysis ± SD of 5 independent assays. \**P*<0.05 compared to the relative control.

Cell fractionation of 8505c cells transiently or stably transfected with PD-1 was used to demonstrate that PD-1 binding to SHP2 enforced the membrane localization of SHP2. Subcellular fractions of membranes (M) or cytosol (C) were obtained from PD-1 overexpressing and from control cells (pFLAG-PD-1 *vs* pFLAG or pCMV3 PD-1 cl 16 *vs* pCMV3). Enrichment of SHP2 levels in the membrane fractions was observed in PD-1 overexpressing cells compared to empty-vector transfected cells. Normalizations of each extract were obtained by using antibodies to transferrin receptor for membrane fraction and α-tubulin for cytosolic extract (**Figure 4C**). In agreement with these observations, immunofluorescence (IF) assay of PD-1 overexpressing TC cells showed a significant increase of SHP2 staining at the plasma membrane in cells overexpressing PD-1 compared to controls (**Figure 4D and Figure Supplement 4B**).

Furthermore, in 8505c cells transfected with PD-1-GFP, we demonstrated by IF that SHP2 and PD-1-GFP co-localize at the plasma membrane (**Figure Supplement 4C**).

### SHP2 dephosphorylates and activates Ras in TC cells

We then searched for the molecular mechanism of Ras activation mediated by the PD-1/SHP2 complex. We first asked whether PD-1 could enhance GRB2 recruitment by SHP2. To this aim, we used pull-down assays with GST-SH2-GRB2 fusion proteins and co-immunoprecipitation assays showing no increased GRB2 binding to SHP2 in PD-1 transfected TC cells compared to controls (**Figure Supplement 4D**). In accordance with these observations, PD-1 enforced expression did not significantly increase SHP2 tyrosine phosphorylation levels (**Figure Supplement 4A**), on which GRB2 binding to SHP2 is dependent, nor changed substantially GRB2 compartmentalization as demonstrated in cell fractionation experiments (**Figure 4C**).

Since the GRB2/SOS complex is not involved in PD-1-mediated Ras activation, we asked whether Ras could be activated by SHP2 through the dephosphorylation of its inhibitory tyrosine residues (*Bunda et al., 2015, Kano et al., 2016*). We evaluated the phosphatase activity of SHP2 and, in parallel, the levels of Ras tyrosine phosphorylation in cells overexpressing or not PD-1. We used a specific SHP2 phosphorylated substrate in the presence of the Malachite Green tracer, a colorimetric method for the detection of free inorganic phosphate (*Bunda et al., 2015*). We observed that SHP2 phosphatase activity was significantly increased in PD-1-versus empty-vector-transfected TC cells (**Figure 4E**). Similar results were obtained in PD-1 stably transfected cells (**not shown**). Consistently with the increased phosphatase activity of SHP2, Ras total phosphorylation levels, in the presence of PD-1, were significantly reduced in TC cells transfected with pCEFL H-Ras AU5 (**Figure Supplement 4E**). To assess whether Ras dephosphorylation occurs in its inhibitory residues 32 and/or 64 (*Bunda et al., 2015*), we used (pan)Ras immunoprecipitation followed by immunoblotting with anti-phospho Y32 (Ras) or Y64 (Ras) antibodies. These experiments demonstrated that PD-1 enforced expression in 8505c cells reduced the Ras phosphorylation levels in the inhibitory tyrosine residues 32 in pCEFL Ras AU5-transfected cells compared to controls (**Figure 4F**). Similar results were obtained in TPC-1 cells (**not shown**). No differences in phosphorylation levels of inhibitory residues 64 were observed (**not shown**).

Taken together, these data indicate that, in TC cells, PD-1 binds SHP2, which in turn dephosphorylates Ras in its inhibitory tyrosine, thus leading to the activation of the MAPK signalling cascade.

### PD-1-induced biologic activities in thyroid cancer cells require the SHP2/BRAF/MEK signalling proteins

To investigate the role of SHP2 in PD-1 functional activity, we treated TC cells, overexpressing or not PD-1, with siRNA targeting SHP2 (siSHP2 – 100 nM) or with a SHP2 allosteric inhibitor that blocks its phosphatase activity (SHP099 – 350 nM) (*Chen et al., 2016*). As shown in **Figure 5A**, siSHP2 was able to significantly reduce SHP2 protein levels compared to scrambled siRNAs (siCTR). By BrdU incorporation assays, we demonstrated that siSHP2 significantly decreased DNA synthesis (**Figure 5B**) in PD-1-, and to a lesser extent in empty vector-transfected, 8505c cells. Consistently, SHP099 inhibitor significantly reduced PD-1-induced DNA synthesis in 8505c cells (**Figure 5C**).

**Figure 5.**
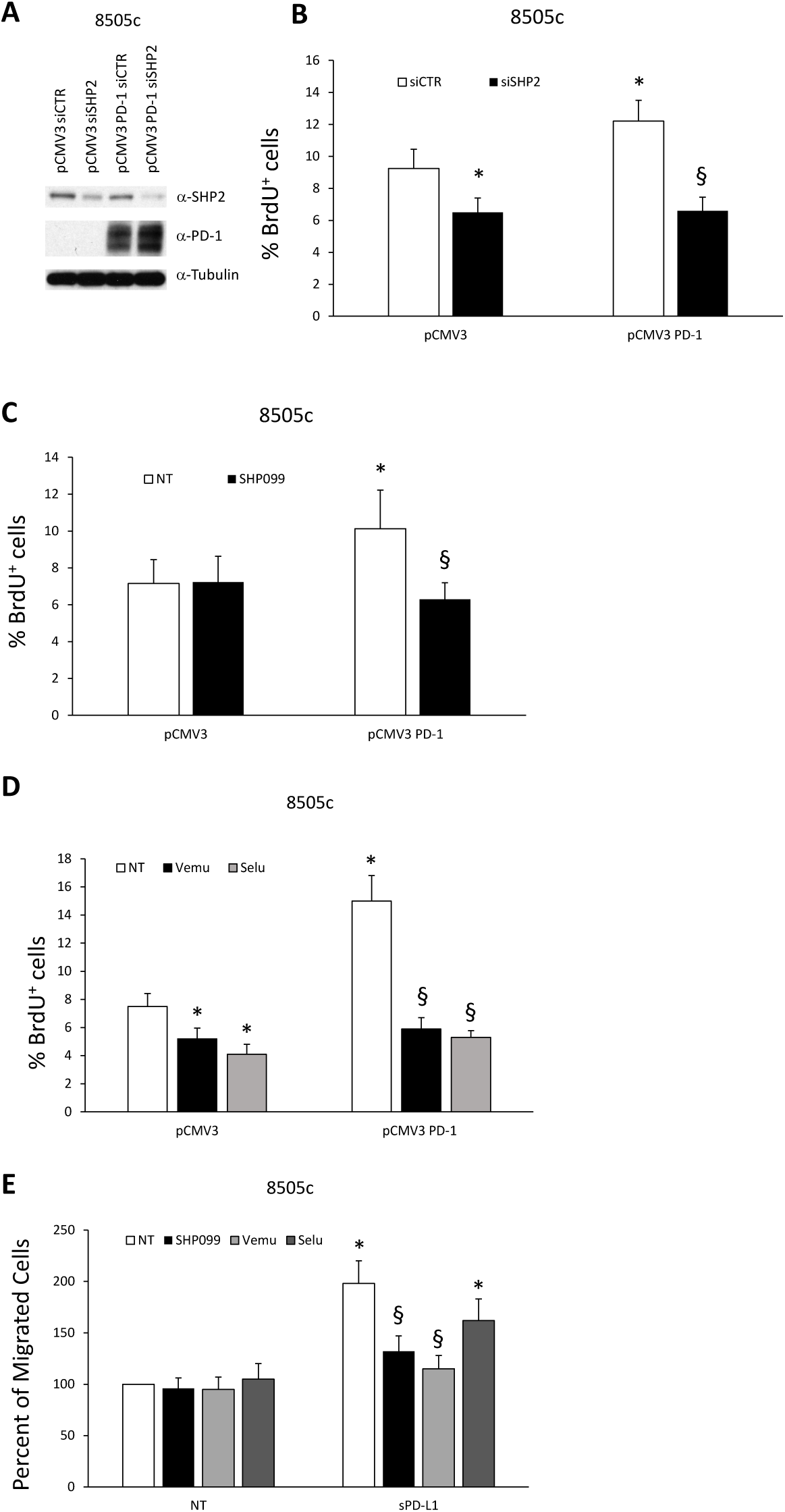
Dependence of PD-1 biologic activities on SHP2/BRAF/MEK cascade. **A**. Effects of siRNA targeting SHP2 (siSHP2 - 100 nM) or the relative control on SHP2 protein expression levels assessed by western blot in 8505c cells stably transfected with PD-1 or the empty vector (one representative clone is shown). **B**. DNA synthesis of stably transfected 8505c pCMV3 PD-1 cells (mean of 3 clones) compared to empty vector transfected cells treated with siSHP2 or siCTR assessed by BrdU incorporation. Data are presented as mean ± SD of 5 independent experiments. **C**. DNA synthesis of 8505c cells stably transfected with PD-1 (pCMV3 PD-1 compared to pCMV3) and treated for 18 h with SHP099 (350 nM) assessed by BrdU incorporation. The mean of three clones is shown for each condition. Data are presented as mean ± SD of 5 independent experiments. **D**. DNA synthesis of stably transfected 8505c pCMV3 PD-1 cells (3 clones) compared to empty vector transfected cells treated for 18 h with Vemurafenib (Vemu – 10 μM) or Selumetinib (Selu - 10 μM) assessed by BrdU incorporation. Data are presented as mean ± SD of 5 independent experiments. **E**. Percent of migrated 8505c cells over control toward sPD-L1 (1 μg/ml) or medium alone (10% FBS) following treatment with SHP099 (350 nM), Vemurafenib (Vemu - 10 μM) or Selumetinib (Selu - 10 μM). Data are presented as mean ± SD of 5 independent experiments. * *P*<0.05 compared to the relative control. § P<0.05 compared to NT or siCTR.

To investigate the role of the downstream signalling cascade in PD-1 dependent biologic TC responses, we conducted BrdU incorporation assays in TC cells overexpressing or not PD-1, in the presence or in the absence of Vemurafenib (Vemu – 10 μM) (*Xing et al., 2011*), a BRAF inhibitor, or Selumetinib (Selu – 10 μM) (*Ball et al., 2007*), a MEK inhibitor. As shown in **Figure 5D**, both drugs were able to significantly revert PD-1-induced DNA synthesis in 8505c cells.

Similar experiments were performed to assess the role of the signalling cascade in PD-1- mediated TC cell migration. **Figure 5E** shows that SHP099 and Vemurafenib, and to a lesser extent Selumetinib, were able to inhibit the migration of 8505c cells induced by sPD-L1. Similar results were obtained in TC cells transfected with PD-1 (**not shown**).

These data demonstrate that PD-1-induced cell proliferation and motility of TC cells are dependent on the SHP2/BRAF/MEK pathway.

### Intrinsic PD-1 signalling enhances xenograft growth of TC cells in immunocompromised mice

To verify whether PD-1 intrinsic signalling and biologic activity could affect tumorigenicity of TC cells, we xenotransplanted 8505c pCMV3 PD-1 (two clones) and control 8505c pCMV3 (a mass population) cells in athymic mice. 8505c pCMV3 PD-1 xenografts displayed increased tumour growth rate that was statistically significant at 4 weeks after injection, in comparison to empty vector transfected cells (**Figure 6A**). End-stage tumours were excised and analysed for cell proliferation (Ki-67), apoptotic rate (cleaved-caspase 3) and vessel density (CD31) by immunohistochemistry. 8505c pCMV3 PD-1 and 8505c pCMV3 xenografts exhibited statistically significant differences in cell proliferation rate, but not in apoptotic rate or vessel density (**Figure 6B** and **Figure Supplement 5A**).

**Figure 6.**
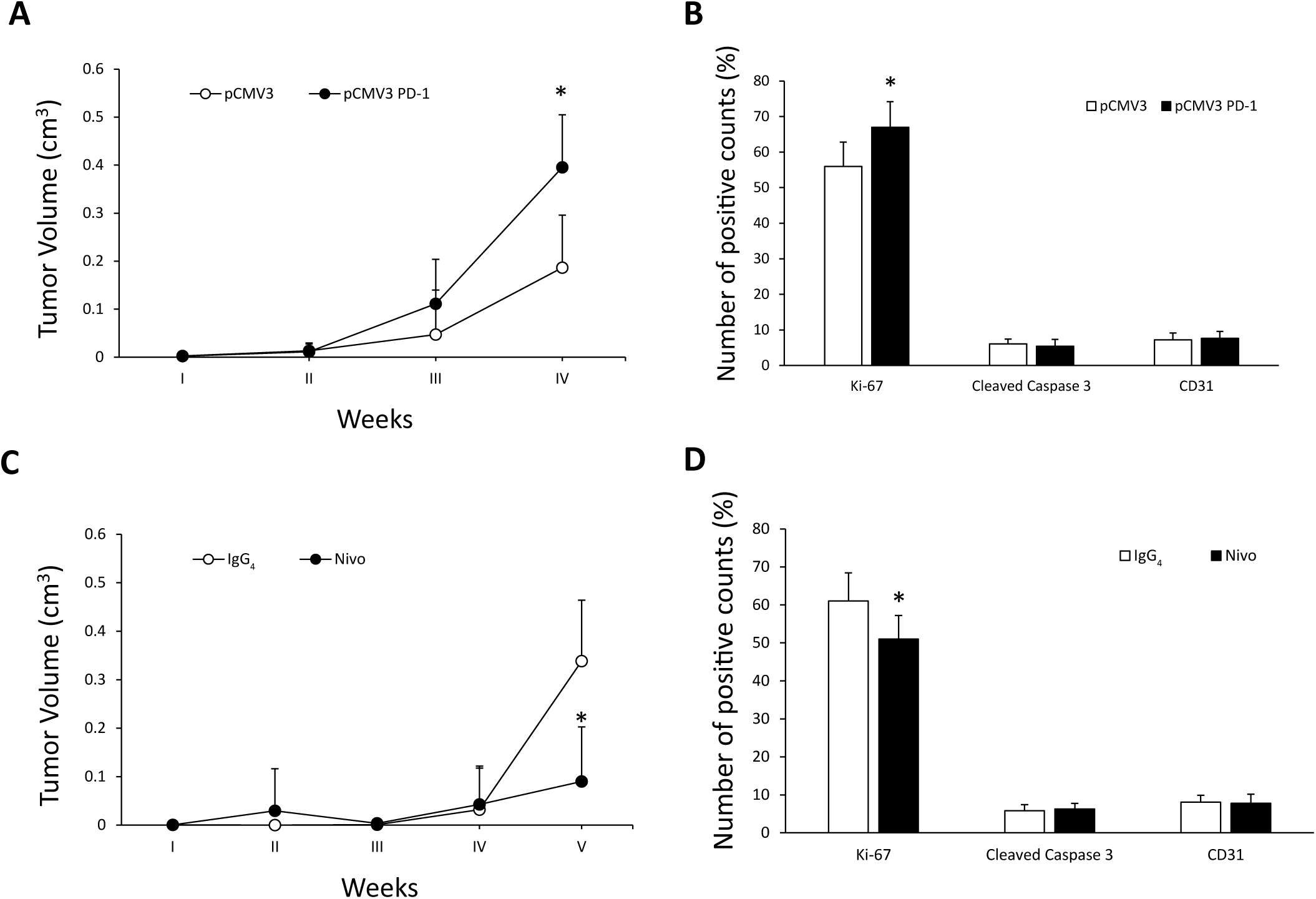
Role of intrinsic PD-1 in tumorigenicity of TC cells. **A**. Tumour growth of pCMV3- (a mass population) or PD-1-transfected (mean of 2 clones) 8505c cells. **B**. Proliferation index (Ki-67), apoptotic rate (cleaved caspase 3), and vessel density (CD31) assessed by immunohistochemistry of 8505c pCMV3 or pCMV3 PD-1 cell xenografts harvested 28 days post-inoculation. The relative quantifications (5 fields/sample) are shown. **C**. Tumour growth of 8505c xenografts in mice treated intraperitoneally (i.p.) at 30▫mg/kg twice per week with Nivolumab (Nivo) or control IgG_4_. **D**. Proliferation index (Ki-67), apoptotic rate (cleaved caspase 3), and vessel density (CD31) assessed by immunohistochemistry of 8505c cell xenografts harvested 35 days post-inoculation. Representative images and the relative quantifications (5 fields/sample) are shown.* P<0.05 compared to the relative control.

To verify whether the inhibition of PD-1 by Nivolumab could affect xenograft growth of parental 8505c cells, mice were xenotransplanted, randomized in two homogeneous groups, and administered intraperitoneally (i.p.) with Nivolumab or control IgG_4_ (30 mg/kg) twice a week. 5 weeks after xenotransplantation, Nivolumab-treated tumours showed a significant decrease in growth rate in comparison with the IgG_4_-treated group (**Figure 6C**). Consistently, Nivolumab significantly reduced TC xenografts’ proliferation without affecting apoptotic rate or vessel density (**Figure 6D** and **Figure Supplement 5B**).

Despite these experiments were carried out in immunocompromised mice, we could not exclude that Nivolumab anti-tumour activity could be ascribed to its ability to affect innate immunity that is present and functional in athymic mice. Thus, we analysed the density and activation of immune cells infiltrating 8505c xenografts treated with Nivolumab or with IgG_4_ by cytofluorimetric analysis. We found that Nivolumab treatment did not change the percentage of CD45+ leucocytes infiltrating xenografts in comparison to IgG_4_ controls, at least at 5 weeks of treatment. Moreover, the density and the expression of polarization/activation markers of tumour-associated macrophages (TAM), of Ly6C+ and Ly6G+ immature myeloid cells, of mature and immature dendritic cells and of regulatory or activated NK, and NKT cells, were comparable between Nivolumab- and IgG_4_-treated 8505c xenografts (**Table Supplement 1**).

These data indicate that, in our model system, PD-1 blockade by Nivolumab inhibits TC cell xenograft growth by affecting tumour cell rather than immune cell compartment.

## Discussion

Several reports point to a promising role of immunotherapy in the treatment of advanced forms of TCs (*Boutros et al., 2016, Saini et al., 2018*). TCGA analysis of TC provided a classification of PTC, in spite of their low mutational burden, as “inflamed” tumours and ATC as hot tumours (*Thorsson et al., 2018*). Interestingly, a profiling of TC confirmed that ATC and PTC are strongly infiltrated by macrophages and CD8^+^ T cells, but that these cells displayed a functionally exhausted appearance (*Giannini et al., 2019*). In TC, high PD-L1 levels significantly correlated with immune infiltrate, increased tumour size and multifocality (*Cunha et al., 2012, Cunha et al., 2013*). Furthermore, the presence of PD-1^+^ T lymphocyte infiltrating TC is associated with lymph-nodal metastasis and recurrence (*French et al., 2012*). Altogether, these data suggest that immune checkpoint inhibitors (ICI) might represent a promising tool for the treatment of these carcinomas.

Our report, for the first time, investigated the expression of the PD-1 receptor in epithelial thyroid cancer cells, demonstrating that a significant percentage of human TC samples displayed PD-1 expression on these cells, although at lower levels compared to the expression found on immune cells infiltrating the tumour. Consistently with the evidence obtained for PD-L1 (*Cunha et al., 2012, Cunha et al., 2013*), our data indicate that PD-1 expression levels correlated with tumour stage and lymph-nodal metastasis in TC. Accordingly, we demonstrated that PD-1 activity could induce proliferation and motility of TC cells in culture. This suggests that the PD-1 intrinsic pathway might have a role in TC cell aggressiveness and invasive ability.

The expression of PD-1 on cancer cells, rather than on immune cells, has been observed recently in melanoma and hepatocellular carcinoma (HCC) (*Kleffel et al., 2015, Li et al., 2017, Yao et al., 2018*). In these cancer types, intrinsic PD-1 activity sustains tumour growth through a ribosomal mTOR/S6K1 signalling (*Kleffel et al., 2015, Li et al., 2017, Yao et al., 2018*). In TC cells, similarly to melanoma and HCC, PD-1 intrinsic signalling sustains cancer cell proliferation, but at variance from these neoplasias, this biologic activity is mediated by the activation of the Ras/MAPK pathway. Interestingly, mutations causing the activation of the Ras/MAPK signalling pathway are found in >70% of PTC (e.g., *RET/PTC* rearrangements and point mutations of the *BRAF* and *Ras* genes) and regulate transcription of key genes involved in TC cell proliferation (*Nikiforov, 2008*). Thus, PD-1 expression could provide a selective advantage to some TC by enhancing the activation of MAPK pathway, thus promoting proliferation and migratory behaviour of cancer cells. Interestingly, besides PD-1, also the immune-checkpoint Cytotoxic T lymphocyte-associated antigen 4 (CTLA-4), classically expressed on leukocytes, has been found to be expressed and functional on cancer cells (*Contardi et al., 2005, Passariello et al., 2020*).

Our data also highlighted the key role of the SHP2 tyrosine-phosphatase in PD-1-mediated tumorigenic activity of TC cells. Interestingly, SHP2 is recruited by PD-1 in T lymphocytes, and inhibits immune receptor signalling by dephosphorylating several downstream substrates (*Rota et al., 2018*). In cancer cells, SHP2 has been described to exhibit oncogenic properties (*Zhang et al., 2015, Ran et al., 2016*). SHP2 functions as an adapter that binds various receptor tyrosine kinases (RTKs) and activates the Ras/MAPK cascade by recruiting the GRB2/SOS complex on the plasma membrane, thus enhancing SOS-mediated GTP loading on Ras (*Zhang et al., 2015, Ran et al., 2016*). SHP2 has also been described as a direct activator of the Ras GTPase through its ability to dephosphorylate specific inhibitory tyrosine residues (*Scott et al., 2011, Bunda et al., 2015, Kano et al., 2016*). In our model system, we found that PD-1 binds SHP2, enhancing its membrane localization and phosphatase activity, thus leading to Ras activation by dephosphorylating inhibitory tyrosines.

Interestingly, increased SHP2 expression was detected in TC samples compared to normal thyroids correlating with poor tumour differentiation, TNM stage and lymph-nodal metastasis (*Hu et al., 2015*). These evidences suggest that SHP2 may represent a potential target for TC therapy both alone and in combination with PD-1 targeting.

Our observations demonstrate that PD-1 is expressed on TC cells and exerts an intrinsic pro-tumorigenic function. Defining the functional and biochemical activity of immune checkpoints both in cancerous cells and in tumour microenvironment of TC will expand our knowledge allowing to develop rational therapeutic strategies for aggressive TC exploiting ICI in combination with SHP2 or kinase inhibitors. In few case reports or in “basket clinical trials” in which ICI [i.e., Pembrolizumab (anti-PD-1), Nivolumab (anti-PD-1), or Atezolizumab (anti-PD-L1)] were used alone or in combination with Multikinase Inhibitors (MKI) for the treatment of advanced and/or metastatic TC, encouraging preliminary clinic evidence of efficacy has been reported (*Cabanillas et al., 2018, Iyer et al., 2018, Liotti et al., 2019*).

The evaluation of PD-1 expression in cancer cells might be important to identify tumours and/or patients that will be likely to respond to ICI administration by taking advantage of both drug effects on immune compartment and on cancer cell proliferation.

## Materials and Methods

### Reagents

pCMV3 and pCMV3 PD-1 plasmids were from Sinobiological (Wayne, PA, USA), pCEFL and pCEFL AU5-tagged Ras (V12) plasmids were a kind gift of J.S. Gutkind (*De Falco et al., 2007*). PD-1 was cloned in pFLAG 5A (Invitrogen, Carlsbad, CA, USA). Soluble PD-L1 (sPD-L1) was from R&D systems (Minneapolis, MN, USA), Nivolumab was kindly provided by S. Scala. Anti-Ras antibody for immunoprecipitation (clone MA1012) was from Invitrogen. Anti-phospho Y32, anti-phospho Y64 Ras antibodies and Y32 and Y64 peptides, used to saturate aspecific binding of each antibody, were provided by M. Ohh. SHP099, Vemurafenib, and Selumetinib were from Selleckchem (Houston, TX, USA). IgG_4_ control antibodies were from Invitrogen.

### Cell culture and transfection

Human thyroid cancer cell lines BcPAP, TPC-1, 8505c, CAL62, SW1736, FRO, BHT101, HTH7 and OCUT1 were obtained and maintained as previously described (*Liotti et al., 2017*). The normal thyroid cells H-6040, isolated from normal human thyroid tissue and cultured in Human Epithelial Cell Medium with the addition of Insulin-Transferrin-Selenium, EGF, Hydrocortisone, L-Glutamine, antibiotic-antimycotic solution, Epithelial Cell supplement, and FBS were purchased from Cell Biologics (Chicago, IL, USA). H-6040 cells were used at passages between 3 and 6.

Transient transfections of TC cells were performed using polyethylenimine according to manufacturer’s instructions (Merck, Darmstadt, Germany). Cells were harvested 48 hrs after transfection. Electroporation was used (Neon® Transfection System for Electroporation, Life Technologies, Carlsbad, CA, USA) to obtain stably transfected cells (*Prevete et al., 2017*).

For RNA interference, we used SMART pools of siRNA from Dharmacon (Lafayette, CO, USA) targeting PD-1 or SHP2. Transfection was carried out by using 100 nM of SMARTpool and 6 μl of DharmaFECT (Dharmacon) for 48 h (*Prevete et al., 2015*).

### Cytofluorimetric analysis

Cells were incubated (30 min at 4°C) with specific or isotype control antibodies. Cells were analysed with a FACS Calibur cytofluorimeter using CellQuest software (BD Biosciences, Mississauga, ON, Canada). 10^4^ events for each sample were acquired (*Prevete et al., 2015*). Anti-PD-1 and anti-PD-L1 antibodies were from ebioscience (Thermo Fisher, Waltham, MA, USA), anti-PD-L2 from Miltenyi Biotec (Bergisch Gladbach, Germany).

### Immunohistochemistry

Thyroid carcinomas were selected from the Pathology Unit of the University of Perugia upon informed consent; the protocol for the study was approved by the institutional committee of University of Perugia. Thyroid tissues were formalin fixed and paraffin embedded (FFPE). Sections of 4 µm were obtained. BOND-III fully automated immunohistochemistry stainer (Leica Biosystems, Wetzlar, Germany) carried out the immunostaining, using heat-induced antigen retrieval at pH 9.0 for 20 minutes, followed by primary antibody (PD-1, clone EH33; dilution 1:200) (Cell Signaling, Beverly, MA, USA) incubation for 15 minutes. Finally, the ready to use Bond(tm) Polymer Refine Detection System allowed the detection of antigen-antibody reaction (*Giannini et al., 2019*). We used a cut-off of 5% to determine the positivity of immunohistochemistry: cases showing immunostaining in more than 5% of neoplastic cells were considered positive, regardless of the intensity of the staining.

### S-phase entry

S-phase entry was evaluated by Bromodeoxyuridine (BrdU) incorporation. Cells were serum-deprived and treated with stimuli for 24 h. BrdU was added at a concentration of 10 μM for the last 1 h. BrdU-positive cells were revealed with Texas Red conjugated secondary Abs (Jackson Laboratories, West Grove, PA, USA). Fluorescence was detected by FACS analysis (*Liotti et al., 2017*).

### Migration assays

Chemotaxis was evaluated using a Boyden chamber. We used a 48-well microchemotaxis chamber (NeuroProbe, Gaithersburg, MD, USA) and 8-μm-pore polycarbonate membranes (Nucleopore, Pleasanton, CA, USA) coated with 10 μg/ml fibronectin (Merck) as described elsewhere (*Prevete et al., 2015*).

### Protein studies

Protein extraction and immunoblotting experiments were performed according to standard procedures (*Collina et al., 2019*). Antibodies to PD-1, phospho-PD-1, phospho-BRAF, phospho-MEK1/2, phospho-MAPK (p44/p42), Ras, phospho-SHP2, SHP2, and GRB2 for Western blot analysis were obtained from Cell Signaling Technology (Danvers, MA, USA). Monoclonal anti-tubulin antibody was from Sigma Aldrich. Secondary anti-mouse and anti-rabbit antibodies were coupled to horseradish peroxidase (Biorad, Hercules, CA, USA).

Cell lysates were subjected to immunoprecipitation with different antibodies or subjected to pull-down binding assays with purified recombinant proteins immobilized on agarose beads. The glutathione-S-transferases (GST) fusion proteins were expressed in *Escherichia coli* and purified with glutathione-conjugated agarose beads (Merck) by standard procedures. The protein complexes were eluted and resolved by sodium dodecyl sulphate-polyacrylamide gel electrophoresis (SDS-PAGE). Immunoblotting with specific antibodies and enhanced chemiluminescence (ECL; Thermo Fisher) were employed for immune-detection of proteins in complexes (*Avilla et al., 2011*).

Cell fractionation experiments were performed using the Subcellular Protein Fractionation Kit for Cultured Cells according to manufacturer’s instructions (Thermo Fisher). Membrane fraction’s protein content was normalized by using anti-transferrin receptor antibody (Invitrogen).

### Immunofluorescence

Cells, grown on coverslips, were washed with phosphate-buffered saline (PBS), fixed with 4% paraformaldehyde (PFA) and quenched with 50 mM NH_4_Cl. Then, cells were permeabilized with 0.2% Triton X-100 for 5 min and blocked for 30 min in PBS containing 5% FBS and 0.5% bovine serum albumin (BSA). Primary antibodies were detected with Alexa Fluor546-conjugated secondary antibodies (Abcam, Cambrige, UK). Images were acquired using a laser scanning confocal microscope (LSM 510; Carl Zeiss MicroImaging, Inc, Oberkochen, Germania.) equipped with a planapo 63X oil-immersion (NA 1.4) objective lens by using the appropriate laser lines and setting the confocal pinhole to one Airy unit. Z-slices from the top to the bottom of the cell by using the same setting (laser power, detector gain) were collected as previously described (*Paladino et al., 2008*).

### SHP2 activity assay

SHP2 phosphatase activity was determined using the human/mouse/rat active DuoSet IC kit (R&D Systems). Briefly, cellular SHP2 bound to anti-SHP2 antibody conjugated to agarose beads was exposed to synthetic phosphopeptide substrate, which is only dephosphorylated by active SHP2 t. The amount of free phosphate generated in the supernatant was determined, as absorbance at 620 nm, by a sensitive dye-binding assay using malachite green and molybdic acid and represents a direct measurement of SHP2 activity in the cellular system (*Bunda et al., 2015*).

### Tumorigenicity in immunocompromised mice

Each group of 8 mice (6-week-old CD1 nu/nu females) was inoculated subcutaneously with 8505c parental cells, 8505c transfected with pCMV3 or pCMV3 PD-1 cells (1×10^7^cells/mouse, two clones) (*Liotti et al., 2017*). Nivolumab (anti-PD-1) or control IgG_4_ were intraperitoneally (i.p.) administered at 30 mg/kg twice per week. The experimental protocol for animal studies was approved by the Ministero Italiano della Salute (No. 317/2019-PR). For xenograft histological analysis, anti-Ki-67 was from Biocare Medical (Pacheco, CA, USA), anti-CD31, anti-cleaved caspase 3 were from R&D Systems.

### Statistical analysis

The results are expressed as the mean ± SEM of at least 3 experiments. Values from groups were compared using the paired Student *t* test or Duncan test. The association between PD-1 expression and clinic-pathologic parameters in immunohistochemistry experiments was conducted using χ2. *P* value < 0.05 was considered statistically significant.

## Acknowledgements

We are grateful to Rosaria Catalano and Mariarosaria Montagna for technical help, and Massimiliano Cacace for animal handling

## Financial supports

Associazione Italiana per la Ricerca sul Cancro (AIRC) IG grant 23218; Istituto Superiore di Oncologia grant (MIUR PON01_02782/12); POR Campania FESR 2014-2020 “SATIN” grant; POR Campania FESR 2014-2020 “RARE.PLAT.NET” grant; and “POR Campania FESR 2014-2020 “grant

## Competing interests

The authors declare no conflict of interest

## Notes

### Competing Interest Statement

The authors have declared no competing interest.

