## Supplementary files for "A cell-autonomous PD-1/PD-L1 circuit promotes tumorigenicity of thyroid cancer cells by activating a SHP2/Ras/MAPK signalling cascade"

**Supplementary data**

**Reagents**

Anti-PD-L1 and anti-PD-L2 blocking antibodies were from R&D systems (Minneapolis, MN, USA); pCMV6 PD-1-GFP from Origene (Rockville, MD, USA), pCMV3 PD-L1 and pCMV3 PD-L2 plasmids were from Sinobiological (Wayne, PA, USA). $IgG_1$ control antibody was from Invitrogen (Carlsbad, CA, USA).

**RNA, cDNA and real-time-PCR**

Total RNA was isolated and retro-transcribed as previously described. Real-time quantitative PCR was performed as reported elsewhere (*Prevete et al., 2017*). Normalization was performed using β-actin and GAPDH mRNA levels. Primer sequences are GAPDH forward 5'-ctgccactgaaaaggaggag-3' and reverse 5'-ttggcactccttgggttatc-3', PD-1 5'-cgcccttgtgctgatcctg-3' and reverse 5'-tgctggtagtggtacatctcc-3', PD-L1 forward 5'-agatgtgaaattgcaggatgcagg-3' and reverse 5'-caattccaagagagaggagaagct-3' and PD-L2 forward 5'-atgatcttcctcctgctaat-3' and reverse 5'-tcagatagcactgttcacttccctc-3'.

**TUNEL Assay**

For the TUNEL assay, an equal number ($5 \times 10^5$) of cells was pleated in 60mm cell culture plates; cells were serum-deprived for 12 h, treated with different stimuli for 24 h and subjected to the TUNEL reaction (Roche, Basel, Switzerland) as described elsewhere (*Prevete et al., 2015*). Fluorescence was detected by FACS analysis.

**Mouse immune cells FACS analysis**

Mice xenografts were dissociated in single cell suspension by using GentleMACS (Miltenyi). For specific immune cell population recognition: anti-CD45, anti-CD11c, and anti-MHC II were used in combination for the detection of immature and mature dendritic cells; anti-CD45, -CD11b, -Ly6C and -Ly6G for the detection of subpopulations of myeloid cells; anti-CD45, -NK1.1, -Cd3ε, -Ly49a, -Ly49c/f, -CD107a for the detection of NK and NKT cells; anti-CD45, -F4/80, -IL12, -IFNγ, IL4, IL-10 for M1 or M2 macrophages. All the antibodies were from Miltenyi Biotec. When necessary, cells were permeabilized using the Cytofix/Cytoperm kit (BD Biosciences). Cells were analysed with a FACS Fortessa using Diva software (BD Biosciences).

| **Cell Population** | **IgG4** | **Nivolumab** | **P** |
| --- | --- | --- | --- |
| Leukocytes ($CD45^+$) | 17.1 ± 4.4 | 17.0 ± 7.7 | 0.97 |
| Dentritic cells | 8.3 ± 2.4 | 7.5 ± 4.1 | 0.62 |
| Immature ($CD45^+$, $CD11c^+$, MHC $II^-$) | 1.2 ± 1.1 | 1.0 ± 0.9 | 0.47 |
| Mature ($CD45^+$, $CD11c^+$, MHC $II^+$) | 5.5 ± 2.2 | 2.0 7.8 ± 4.1 | 0.15 |
| Macrophage | 35.5 ± 5.8 | 30.3 ± 4.9 | 0.19 |
| M1 ($CD45^+$, $F4/80^+$, $IL12^+$, $IFN\gamma^+$) | 5.9 ± 6.1 | 2.7 ± 1.7 | 0.43 |
| M2 ($CD45^+$, $F4/80^+$, $IL10^+$, $IL4^+$) | 3.8 ± 1.4 | 4.4 ± 1.6 | 0.52 |
| Myeloid cells | 5.9 ± 2.1 | 6.7 ± 5.6 | 0.82 |
| Ly6C ($CD45^+$, $CD11b^+$, $Ly6C^+$, $Ly6G^-$) | 0.6 ± 0.2 | 1.1 ± 0.9 | 0.47 |
| Ly6G ($CD45^+$, $CD11b^+$, $Ly6C^-$, $Ly6G^+$) | 6.2 ± 0.9 | 8.5 ± 6.6 | 0.49 |
| NK-T cells ($CD45^+$, $NK1.1^+$, $Cd3\varepsilon^+$) | 3.1 ± 1.3 | 2.8 ± 2.6 | 0.85 |
| NK cells | 0.4 ± 0.3 | 1.7 ± 2.5 | 0.33 |
| Regulatory ($CD45^+$, $NK1.1^+$, $Cd3\varepsilon^-$, $Ly49a^-$, $Ly49c/f^-$, $CD107a^+$) | 34.8 ± 6.2 | 43.3 ± 14.4 | 0.44 |
| Killing ($CD45^+$, $NK1.1^+$, $Cd3\varepsilon^-$, $Ly49a^+$, $Ly49c/f^+$, $CD107a^-$) | 41.0 ± 7.8 | 43.8 ± 17.0 | 0.81 |

**Table Supplement 1.** Mouse immune cell density (expressed as percentage of $CD45^+$ leukocytes) in 8505c xenografts.

### Supplementary Figure Legend

**Figure Supplement 1 Immune checkpoint expression in thyroid cancer (TC) cells.**

mRNA expression indicated as ΔCt for PD-1, PD-L1 and PD-L2 in H-6040 normal thyroid epithelial cells, PTC-derived cell lines (BcPAP and TPC-1), and ATC-derived cell lines (8505c, CAL62, SW1736, FRO, BHT101, HTH7, OCUT1). Data are presented as mean ± SD of 5 independent experiments.

**Figure Supplement 2 Functional activity of intrinsic PD-1 circuit in TC cells.**

**A.** Expression levels of PD-1 in 8505c and TPC1 cells or in 8505c and TPC-1 transiently transfected with pFLAG or pFLAG PD-1, assessed by western blot. **B.** Cell cycle distribution of 8505c and TPC-1 cells transiently transfected with pFLAG or pFLAG PD-1, measured by Propidium Iodide (PI) staining by means of Flow Cytometry. The percent of the cells distributed in G0/G1, S, G2/M was indicated in each panel. Representative experiments are shown. **C.** Percent of apoptotic cells assessed by TUNEL reaction in 8505c and TPC-1 cells transiently transfected with pFLAG or pFLAG PD-1 and treated or not with soluble PD-L1 (sPD-L1 - 1 μg/ml). Data are presented as mean ± SD of 5 independent experiments. **D.** Cytofluorimetric evaluation of PD-1 expression in 8505c cells treated with siPD-1 (solid lines) or scrambled siCTR (dotted line) (100 nM). **E.** Cell cycle distribution of 8505c and TPC-1 cells treated with Nivolumab (Nivo - 10 μg/ml) or control $IgG_4$ (10 μg/ml), measured by Propidium Iodide (PI) staining by means of Flow Cytometry. The percent of the cells distributed in G0/G1, S, G2/M was indicated in each panel. Representative experiments are shown. **F.** Percent of apoptotic cells assessed by TUNEL reaction in 8505c and TPC-1 cells treated with siPD-1 (100 nM) or Nivolumab (Nivo - 10 μg/ml) or the relative controls. Data are presented as mean ± SD of 5 independent experiments. **G.** DNA synthesis of 8505c cells transiently transfected with pCMV3, pCMV3 PD-L1 or pCMV3 PD-L2 or treated with anti-PD-L1, anti-PD-L2 blocking antibodies or $IgG_1$ isotype control (10 μg/ml) assessed by

BrdU incorporation. Data are presented as mean ± SD of 5 independent experiments. * P<0.05 compared to the relative control.

**Figure Supplement 3 Signalling pathways downstream PD-1overexpression.**

**A.** Expression levels of PD-1 in some clones or mass population obtained from 8505c cells stably transfection with PD-1, assessed by western blot. **B**. Expression levels of phosphorylated forms of BRAF, MEK1/2 and MAPK (p44/p42) in 8505c cells stably transfected with PD-1 or the empty vector, assessed by western blot. **C**. Activation of AKT, SRC, S6, S6K, 4EBP1 in 8505c and TPC-1 cells, transiently transfected or not with PD-1 or the relative empty vector, assessed by western blot for their phosphorylated forms.

**Figure Supplement 4 Effects of intrinsic PD-1 on SHP2 localization and functions.**

**A.** Expression levels of phospho-PD-1, SHP2 and phospho-SHP2 in 8505c and TPC-1 cells transiently transfected with pFLAG PD-1 or the empty vector pFLAG, assessed by western blot. **B.** Immunofluorescence microscopy of 8505c cells, transiently transfected with pFLAG PD-1 or the empty vector, with antibody specific for SHP2. Bars, 5 μm. **C.** Immunofluorescence microscopy of 8505c cells transiently transfected with pCMV6 PD-1-GFP and stained with antibody specific for SHP2, and the merged signal. Bars, 5 μm. **D.** Total protein extracts from TPC-1 cells transiently transfected with pFLAG-PD-1 or the empty vector pFLAG were subjected to a pull-down assay using the indicated recombinant proteins or to immunoprecipitation using the indicated antibodies. Proteins were immunoblotted with antibody against SHP2 or GRB2. **E**. Total cell protein extracts from 8505c cells transiently transfected with combination of pCEFL H-Ras AU5, pFLAG PD-1 or empty vector (pFLAG + pCEFL) were subjected to immunoprecipitation with anti-phospho tyrosine followed by western blotting with pan(RAS) antibody. A representative experiment is shown, together with the mean densitometric analysis ± SD of 5 independent assays. * P<0.05 compared to the relative control.

**Figure Supplement 5 Immunohistochemical evaluation of 8505c xenografts.**

**A.** Proliferation index (Ki-67) assessed by immunohistochemistry of 8505c pCMV3 and pCMV3 PD-1 cl13 xenografts harvested 28 days post-inoculation. Representative images are shown. **B.** Proliferation index (Ki-67) assessed by immunohistochemistry of 8505c xenografts harvested 35 days post-inoculation in mice treated with Nivolumab or control $IgG_4$. Representative images are shown.

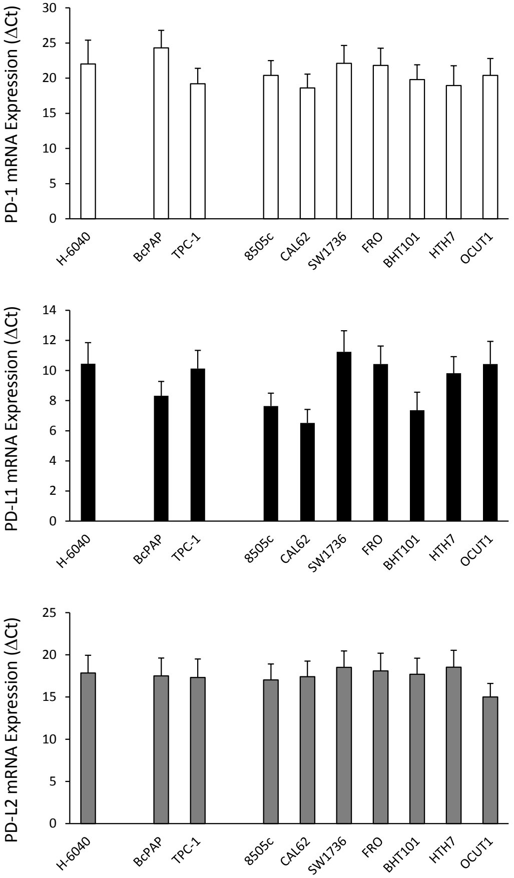


Suppl. Figure 1

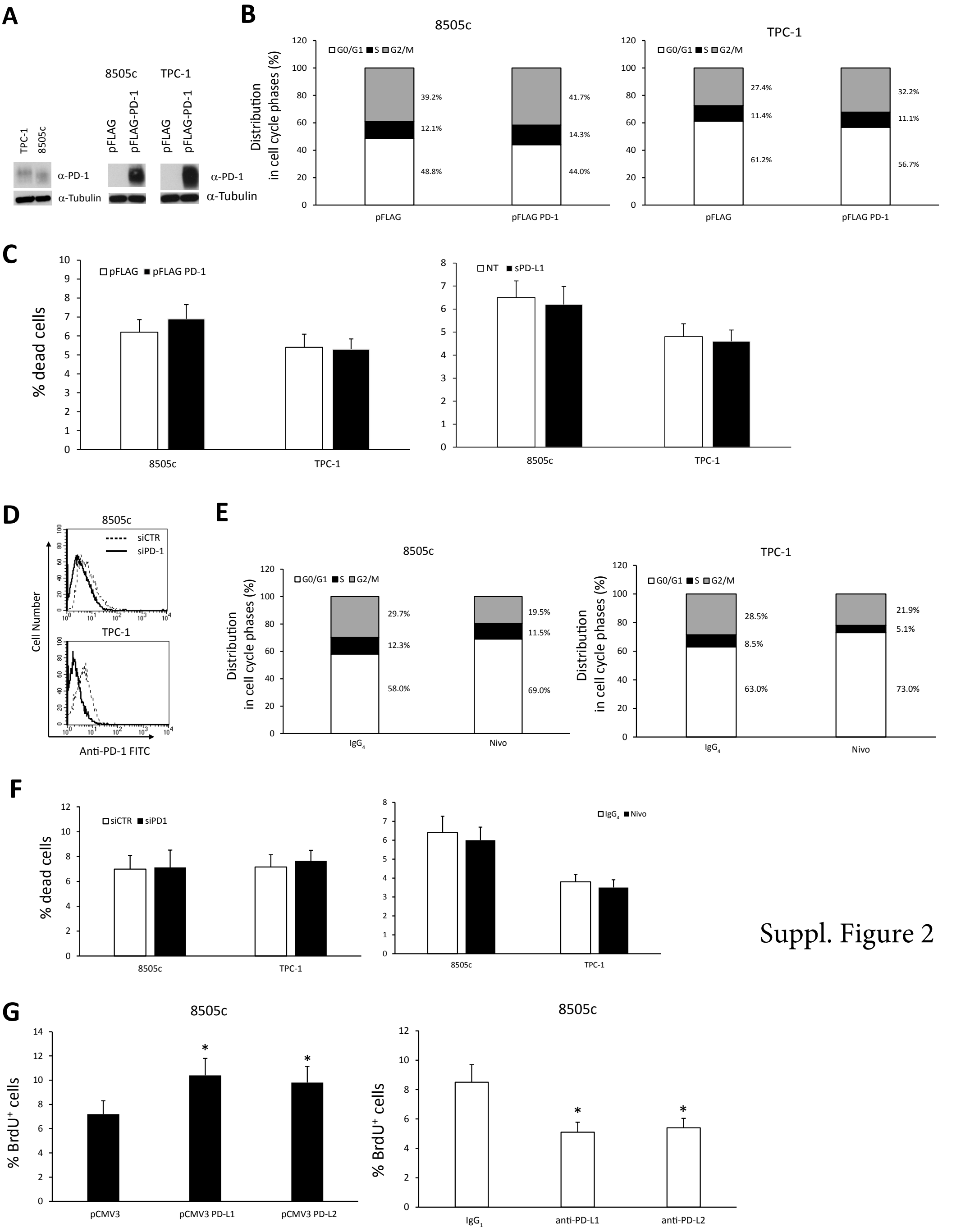


Suppl. Figure 2

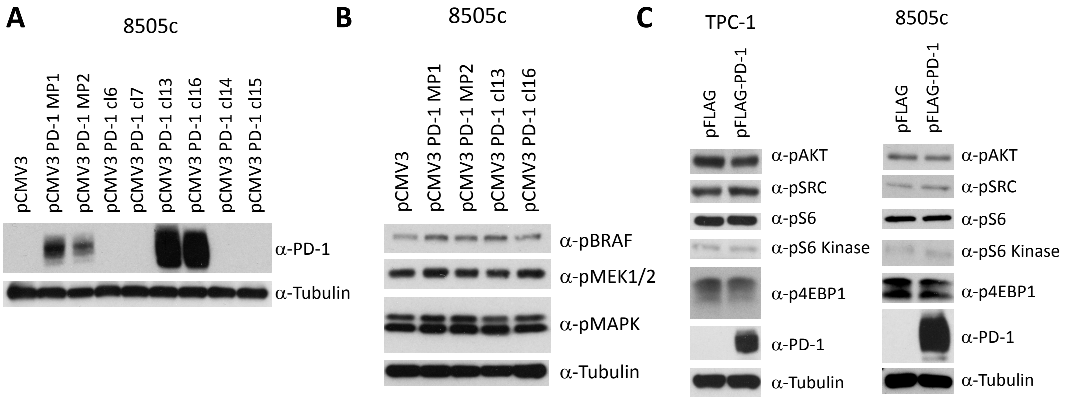


Suppl. Figure 3

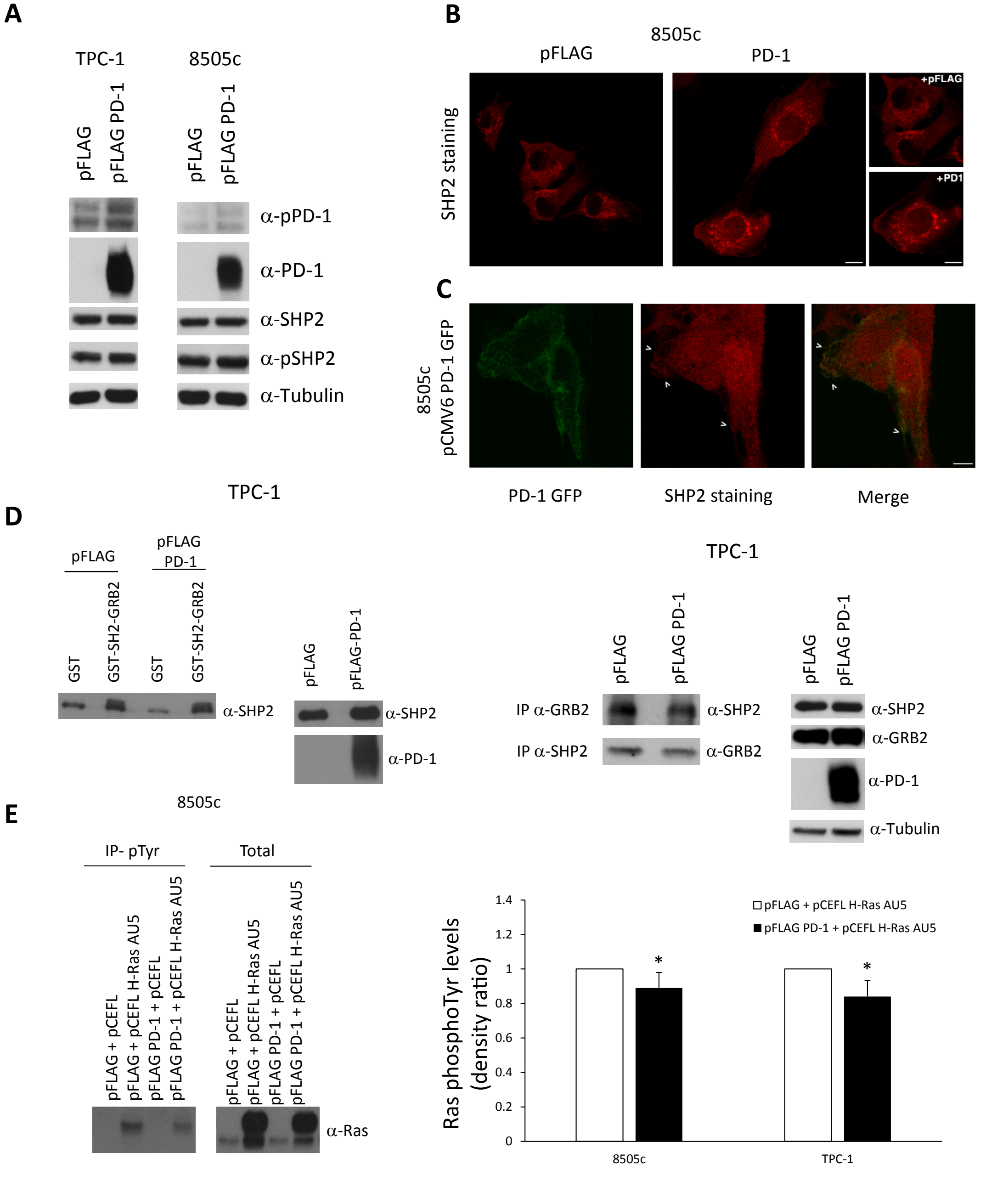


Suppl. Figure 4

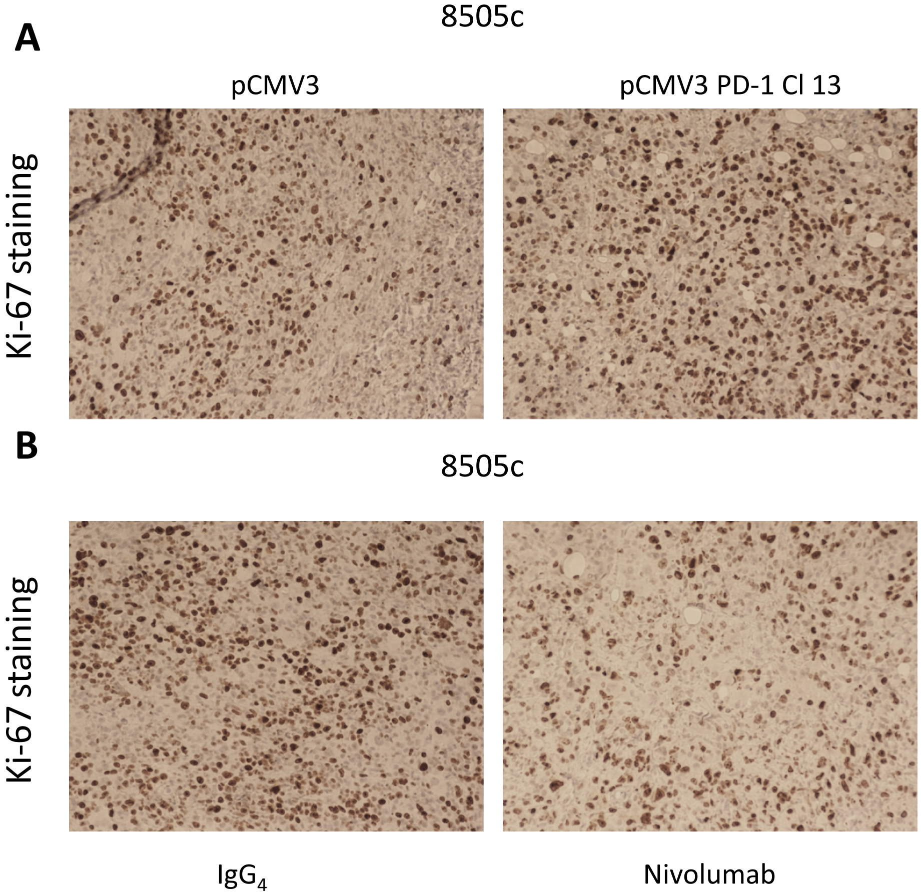


Suppl. Figure 5
